# Longitudinal analysis of the faecal microbiome in pigs fed *Cyberlindnera jadinii* yeast as a protein source during the weanling period followed by a rapeseed- and faba bean-based grower-finisher diet

**DOI:** 10.1101/2021.02.11.430725

**Authors:** S. Iakhno, F. Delogu, Ö.C.O. Umu, N.P. Kjos, I.M. Håkenåsen, L.T. Mydland, M. Øverland, H. Sørum

## Abstract

The porcine gut microbiome is closely connected to diet and is central to animal health and growth. The gut microbiota composition in relation to *Cyberlindnera jadinii* yeast as a protein source in a weanling diet was studied previously. Also, there is a mounting body of knowledge regarding the porcine gut microbiome composition in response to the use of rapeseed (*Brassica napus* subsp. *napus*) meal, and faba beans (*Vicia faba*) as protein sources during the growing/finishing period. However, there is limited data on how the porcine gut microbiome respond to a combination of *C. jadinii* yeast in the weanling phase and rapeseed meal and faba beans in the growing/finishing phase. This work investigated how the porcine faecal microbiome was changing in response to a novel yeast diet with a high inclusion of yeast proteins (40% of crude protein) in a weanling diet followed by a diet based on rapeseed meal and faba beans during the growing/finishing period. The feacal microbiomes of the weanling pigs fed yeast were more diverse with higher relative abundance of *Firmicutes* over *Bacteroidetes* compared with those of soybean meal-based diet fed weanlings. Reduced numbers of *Prevotella* in the yeast fed faecal microbiomes remained a microbiome characteristic up until two weeks after the yeast diet was changed to the rapeseed/faba bean growing finishing diet. A number of differentially abundant bacterial phylotypes along with distinct co-occurrence patterns observed during the growing/finishing period indicated the presence of a “carry-over” effect of the yeast weanling diet onto the faecal microbiomes of the grower/finisher pigs.

## Introduction

Soybean (*Glycine max*) meal (SBM) is a commonly used protein source in commercial livestock diets in Europe. This leads to intensified crop production, which puts pressure on land and water resources, and it reduces their availability as food for humans (reviewed in [1]). Yeast proteins, or yeast-derived nutrients proved a potent alternative to the soybean-based and other conventional protein sources in the feed for weanling piglets [2–4]. In growing/finishing (G/F) pig diets, rapeseed (*Brassica napus* subsp. *napus*) meal (RSM) based formulations are believed to offer the proteins required for animal growth along with the potential of prebiotic properties of RSM which are important for animal health [5, 6]. Gut bacterial consortia play a chief role in the large intestine carbohydrate fermentation whereby supplying the host the molecules valuable for the health and development (e.g. short-chain fatty acids) [7–9]. It has been shown that the replacement of the conventional proteins in weanling pig diets by those derived from yeast has both an impact on the large intestine bacterial composition [10] and positive effects on the pig immune system [11–13]. We previously characterised the compositional changes of the large intestine microbiota in weanling piglets fed *C. jadinii* yeast-based diet. Those changes featured lower alpha microbial diversity in the caecum and colon of the yeast group compared with those of the control group. *Prevotella*, *Mitsuokella* and *Selenomonas* affiliated taxa were more predominant in the yeast associated large intestine microbiomes compared with those of the controls [10]. Umu et al. showed that RSM-based diets during G/F period modulated the porcine gut microbiota favouring the microbial taxa that are linked to an improved gut health state. *Mucispirillum* in the ileum, as well as *Bulleidia*, *Eubacterium*, *Lachnospira*, and *Paraprevotella* in the large intestine, were differentially abundant in the RSM-based G/F pigs (aged 88 days) compared with those of the SBM-fed pigs [14].

While the effects of the SBM diet on the pig gut microbiota were studied separately for weaning period and for G/F period, there is a gap in knowledge on how the porcine gut microbiota respond to a combination of diets wherein the conventional proteins are replaced by the yeast-derived proteins during weaning followed by RSM-based diets during the G/F period. Furthermore, it is not clear whether the yeast diets at weaning have a “carry-over” effect on the pig gut microbiota of the G/F pigs, i.e. the microbiota composition changes due to the yeast diet remain in the G/F period.

To address these questions, we designed a longitudinal study of the porcine faecal microbiomes by using 16S *rRNA* gene metabarcoding sequencing. We characterized the faecal microbiome structure of pigs fed yeast-based weanling diet followed by the RSM-based diet during G/F period (YL group) contrasting it with those fed SBM-based weanling diet followed by the RSM-based diet during G/F period (CL group).

## Results

### The impact of *C. jadinii* yeast proteins on the faecal microbiome bacterial diversity

All animals were healthy during the time of the experiment. There were no major differences in zootechnical performance parameters between the pigs fed the SBM or the RSM diets (S1 Table). Profiling of faecal microbial communities was performed on 8 randomly selected piglets from either YL or CL groups at the following time points: d0 (weaning), d8, and d22 post-weaning (PW). The sampling continued after the introduction of the animals to the grower-finisher RSM-based diet at day 28 PW. The YL and CL faecal samples were collected at d36, d57 and d87 PW. After filtering, denoising, and chimera removal with the DADA2 pipeline, there were on average 62472 (SD=16512) reads per sample available for downstream analyses (S1 Figure). The reads were demultiplexed into 3721 amplicon sequence variants (ASVs) representing the faecal microbiome at d0 (805 ASVs), d8 (1466 ASVs), d22 (2024 ASVs), d36 (1880 ASVs), d57 (2010 ASVs), and d87 PW (2050 ASVs) both feeding groups concerned.

### Alpha diversity

Alpha diversity of the faecal microbiomes in both arms of the study increased between weaning (Shannon index mean = 4.16 (SD=0.38)) and d22 PW (Shannon index mean = 5.07 (SD = 0.24))(Figure 2). When pigs were allocated to the G/F diet, there was a less pronounced increase in the Shannon index of the faecal microbiomes in both arms of the study compared to that of the weaning period. It ranged from 5.07 (SD = 0.24) at d36 PW to 5.35 (SD=0.22) at d87 PW. The pairwise comparison of alpha diversity between the YL and CL groups was accomplished by using DivNet statistical procedure. There was no statistically significant difference in microbial alpha diversity between the CL and YL piglets at the baseline (p=0.69). The Shannon diversity index was higher in YL microbiomes than in the CL ones at d8, d22, and d57 PW, while that was opposite at d36, and d57 PW (Figure 2).

**Figure 1.**
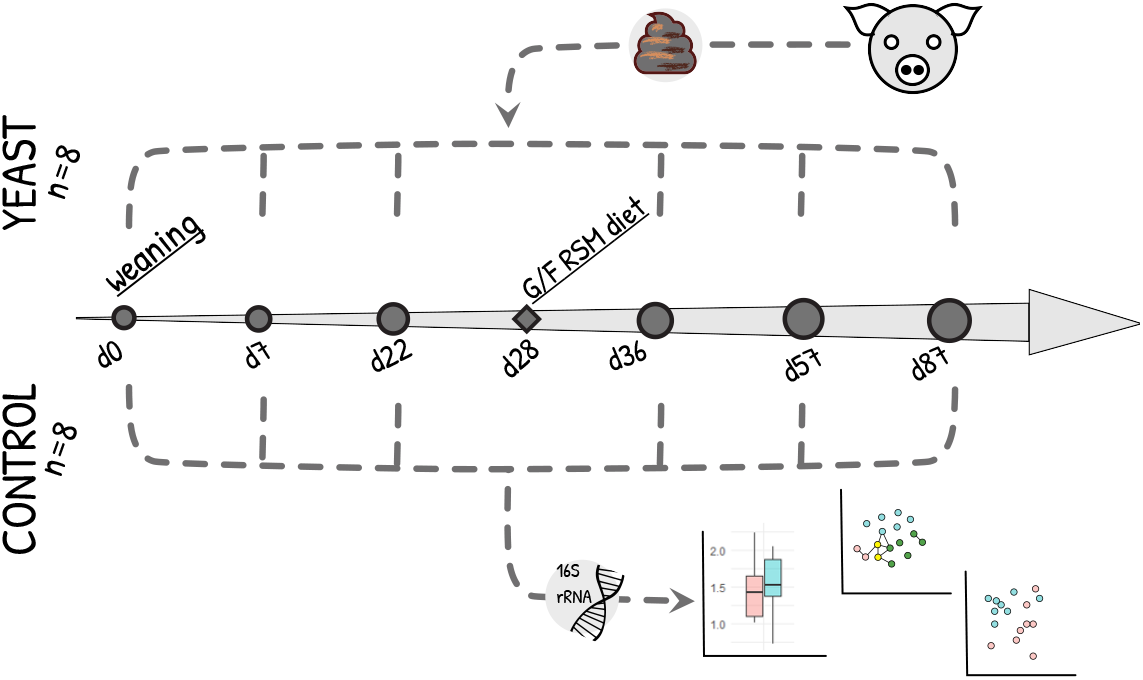
Overview of the experimental design. The timeline of the experiment is shown for two groups of animals in the experiment: YEAST (YL in the text) and CONTROL (CL in the text). Metabarcoding sequencing was done for the faecal samples collected at the days drawn as grey circles (d0, d8, d22, d36, d57, and d87 post-weaning)

**Figure 2.**
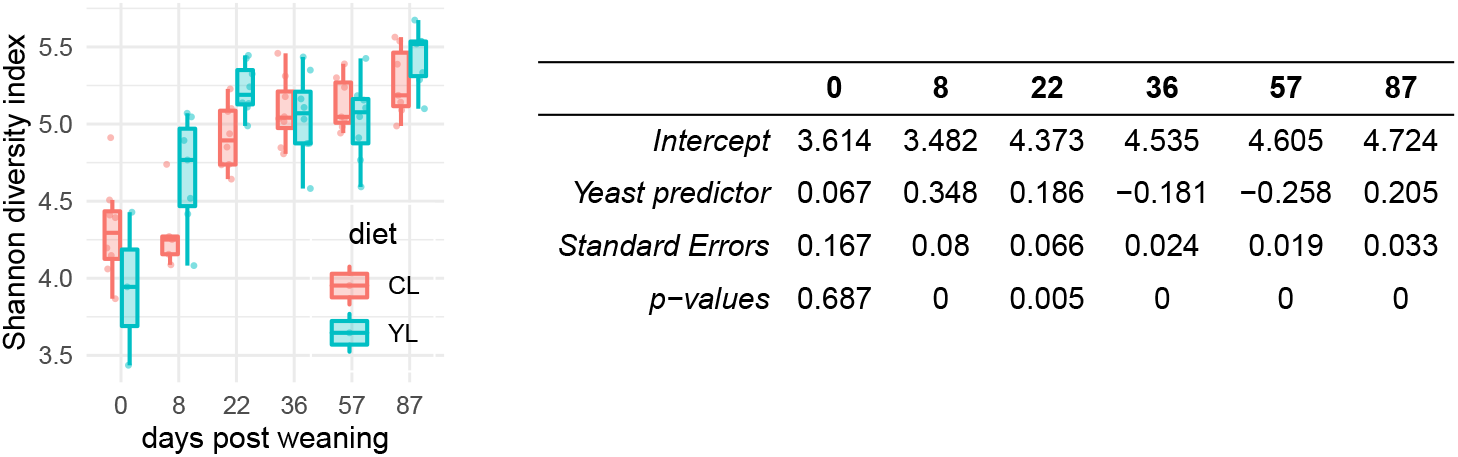
Alpha microbial diversity. **Left** Distributions of the observed values of Shannon diversity index. **Right** The summary of the statistical inference for the alpha diversity measured by Shannon diversity index. The ‘intercept’ terms are the inferred estimates of the control group (CL) Shannon indices across d0-d87 PW. The ‘Yeast predictor’ terms are the inferred estimates of the yeast group (YL) Shannon indices across d0-d87 PW in relation to the ‘intercept’.

### Beta diversity

Beta diversity between the YL and CL pig faecal microbiomes was compared using weighted, and unweighted UniFrac distances as response variables for permutational multivariate analysis of variance (PERMANOVA) test (Figure 3, S2 Table). At the baseline (d0), the microbial communities did not differ, however, at day 8 PW, the diet was predictive of variance in faecal microbiome compositions estimated by UniFrac (F=1.17, R^2^=14.6%, p=0.024) but not for weighted UniFrac (F=0.92, p=0.08). Notably, when “pen” variable was added to the model with the unweighted UniFrac distance as a response term, the prediction of variance in the beta-diversity metric increased to 46.4% (F=1.9, R^2^=14.6%, p=0.007 and F=1.38, R^2^=31.8, p=0.032 for ‘diet’ and ‘pen’ variables respectively). The variance in the weighted UniFrac distance could be predicted by the ‘sow’ variable for d8 PW microbiomes (F=2.03, R^2^=53.6%, P=0.011). At day 22 PW, the diet could predict up to 24.8% of variance in weighted UniFrac distances of the faecal microbiomes (F=4.6, p=0.005) whilst the variance in unweighted UniFrac distance matrices could not be resolved by the diet (p>0.05). There was no difference in beta diversity metrics between the YL and CL pig microbiomes at d36, and d57 PW both distance matrices concerned. Of note, despite there was no effect of diet regimens during grower-finisher period (d36 - d87 PW), the results of PERMANOVA test showed that litter could be predictive of the faecal microbial composition structure. As much as 50.1% of the variance in the faecal microbiomes (weighted UniFrac distance) could be explained by the litter and diet (F=2.33, p=0.033).

**Figure 3.**
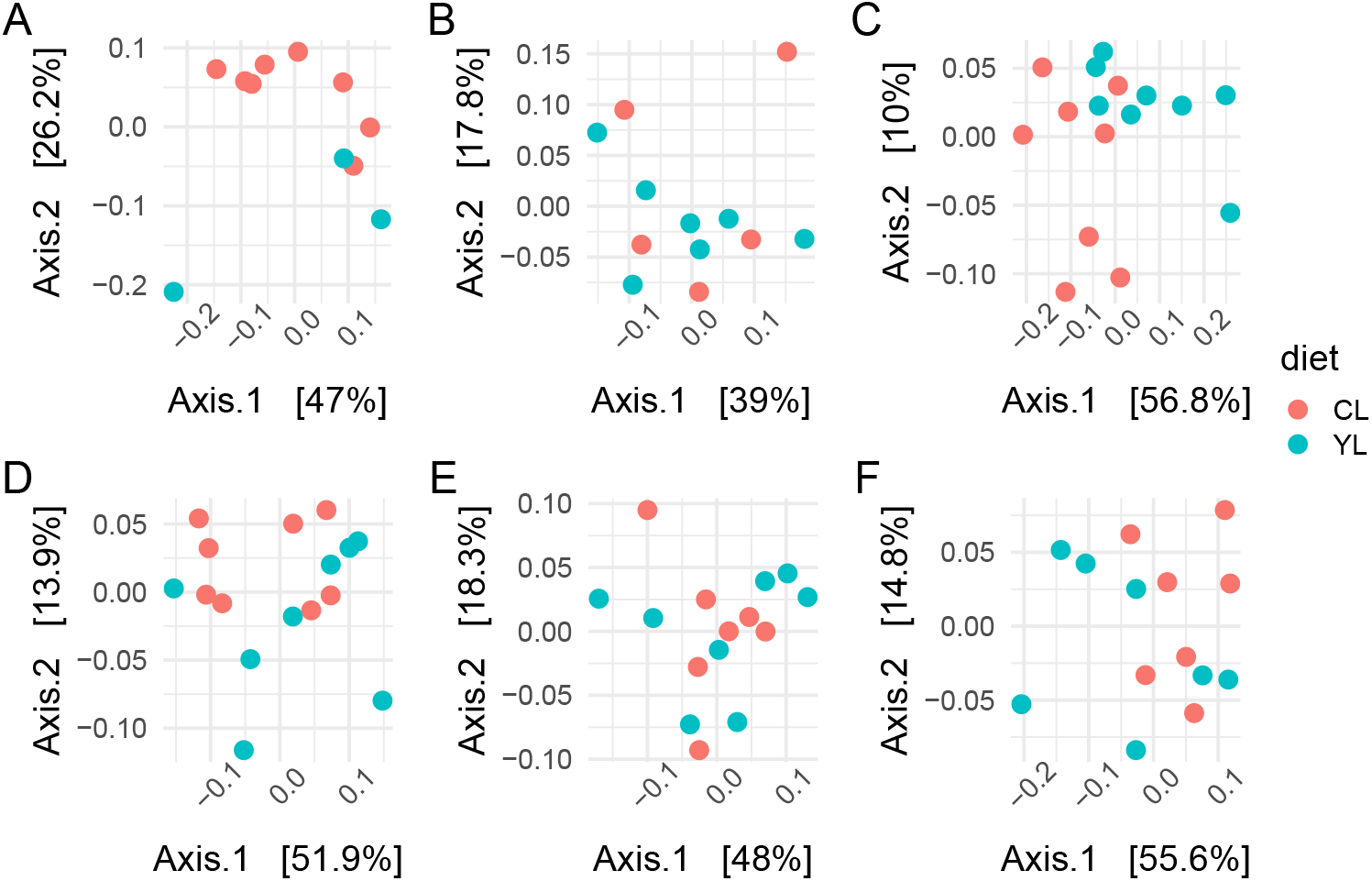
Beta microbial diversity. Principal coordinate analysis plots of the pig faecal microbiomes coloured by diet at day 0 (panel **A**), 8 (panel **B**), 22 (panel **C**), 36 (panel **D**), 57 (panel **E**), and 87 (panel **F**) post-weaning

### Relative abundance of bacterial phylotypes and differential abundance test

Two major bacterial phyla, *Bacteroidetes* and *Firmicutes*, constituted more than 85% of the faecal microbiomes in both feeding groups at all time points with 65.6% (SD = 6.8%) and 24.3% (SD = 4.46%) on average, respectively.

#### Weaning period

At d22 PW, the yeast faecal microbiomes had higher relative abundance of *Firmicutes* (est=0.49, p=0.004) and lower relative abundance of *Bacteroidetes* (est =-0.58, t=-3.81, p=0.002) than those of the SBM-based ones (S2 Figure). Also, for both phyla, *Firmicutes* and *Bacteroidetes*, the variability was lower in the yeast faecal microbiomes compared to those of the SBM-based ones (est =−4.19, t = −11.9, p= 2.31e-08; and est = −3.865, t= −11.03, p= 5.74e-08, respectively). We followed the two differentially abundant and variable phyla up to the class taxonomic level. These were of the *Bacteroidales* and *Clostridiales* orders. The major differences between the faecal microbiomes of YL and CL occurred on d8 PW when species agglomeration was applied (see methods). *Paraprevotellaceae*, *Desulfovibrionacea* ASVs, *Paludibacter*, *Prevotella stercorea*, and *Phascolarcobacterium* ASVs were more predominant in the CL faecal microbiomes at d8 PW compared with those of the YL (Figure 4). Unclassified *Bacteroides*, *Blautia*, unclassified *Ruminococcus*, *R. bromii*, *Sphaerochaeta*, *Treponema*, and *Succiniclasticum* ASVs were differentially abundant in the YL faecal microbiomes at d8 PW. At d22 PW, there were more *Fibrobacter* and *Prevotella*(ASV2) ASVs in the CL faecal microbiomes while *R. bromii* ASV was more abundant in the YL microbiomes (Figure 4).

**Figure 4.**
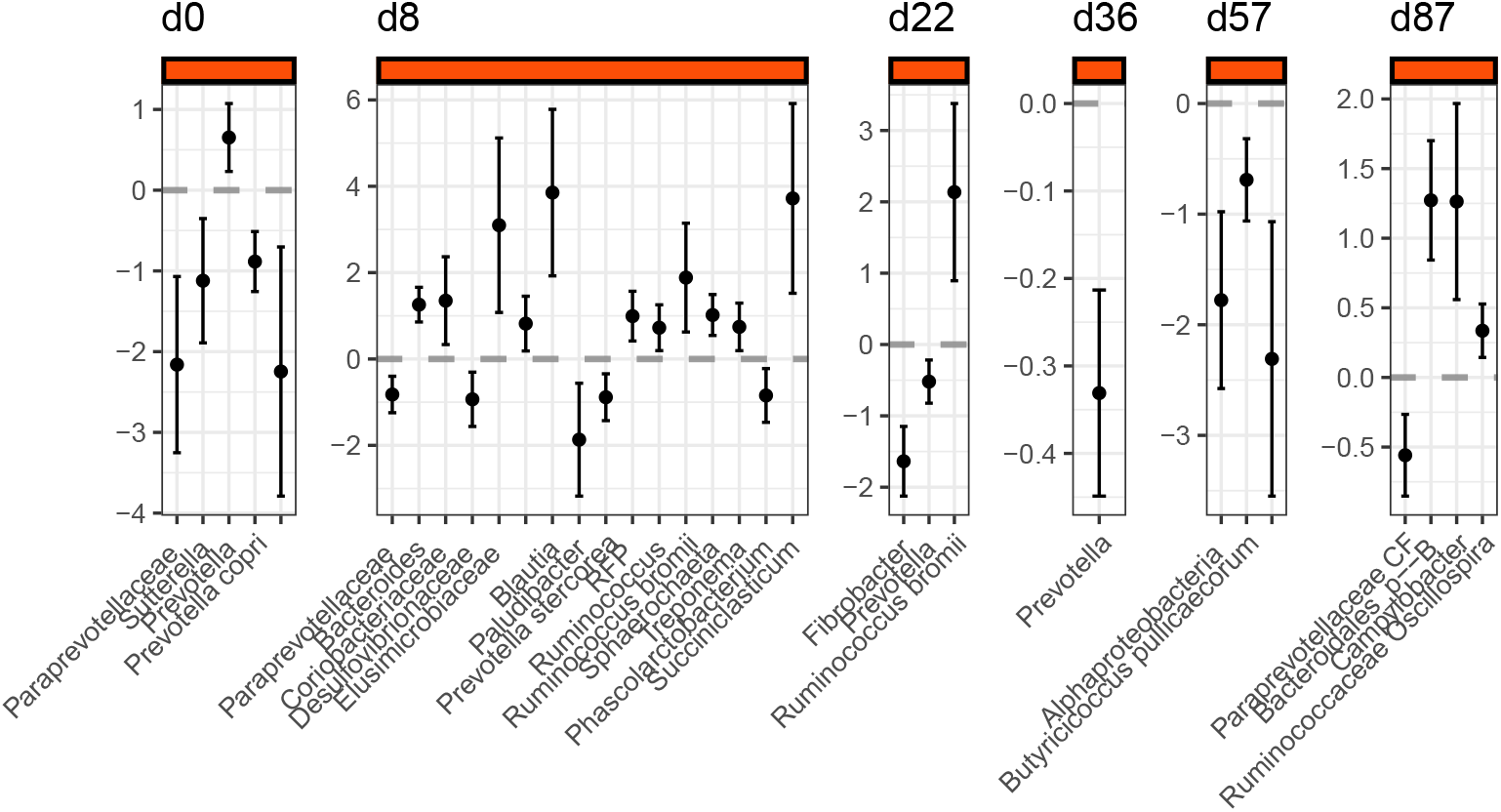
Differentially abundant taxa. The estimates of the beta-binomial regression on the porcine faecal microbiomes along with its standard errors across d0-d87 PW; the positive estimates (above the grey dashed line, “0”) indicate the taxa that are more predominant in the yeast group (YL in the text) compared with those of the controls (CL). Some differentially abundant ASVs are not printed on the X-axis due to taxonomic ambiguity

#### G/F period

At d36 PW, the relative abundance of the same as at d22 PW *Prevotella* (ASV2) was more prevalent in the CL faecal microbiomes (est = −0.33, t=-5.5, p= 0.0004) compared with those of YL diet (Figure 4). At d57 PW, two ASVs, RF32, manually reclassified (see methods) as *Novispirillum* sp., (*Alphaproteobacteria*) and *Butyricicoccus pullicaecorum* were observed at lower relative abundances in the YL faecal microbiomes compared with those of the CL (Figure 4). At d87 PW, *Campylobacter*, *Bacteroidales* order, and *Oscillospira* ASVs relative abundance was higher in the YL faecal microbiomes compared with that of the CL (Figure 4). A *Paraprevotellaceae* ASV was more predominant in the CL faecal microbiomes than those of YL on d87 PW (Figure 4).

### Microbial network analysis

Next, we applied Sparse Inverse Covariance Estimation for Ecological Association Inference approach (SPIEC-EASI) to investigate networked microbial communities’ patterns of the faecal microbiomes of the YL and CL pigs. The connectivity of the networks, i.e. the way the nodes are connected via edges, was sparse and increased moderately over the time; however, no evident difference was present in the two conditions (Figure 5). Moreover, in all the samples the majority of the nodes remained disconnected from the few connected components. Within those, we looked which ASVs (genus level) transited across the CL or YL microbiomes networks consecutively from one point of time to the next (Figure 5).

**Figure 5.**
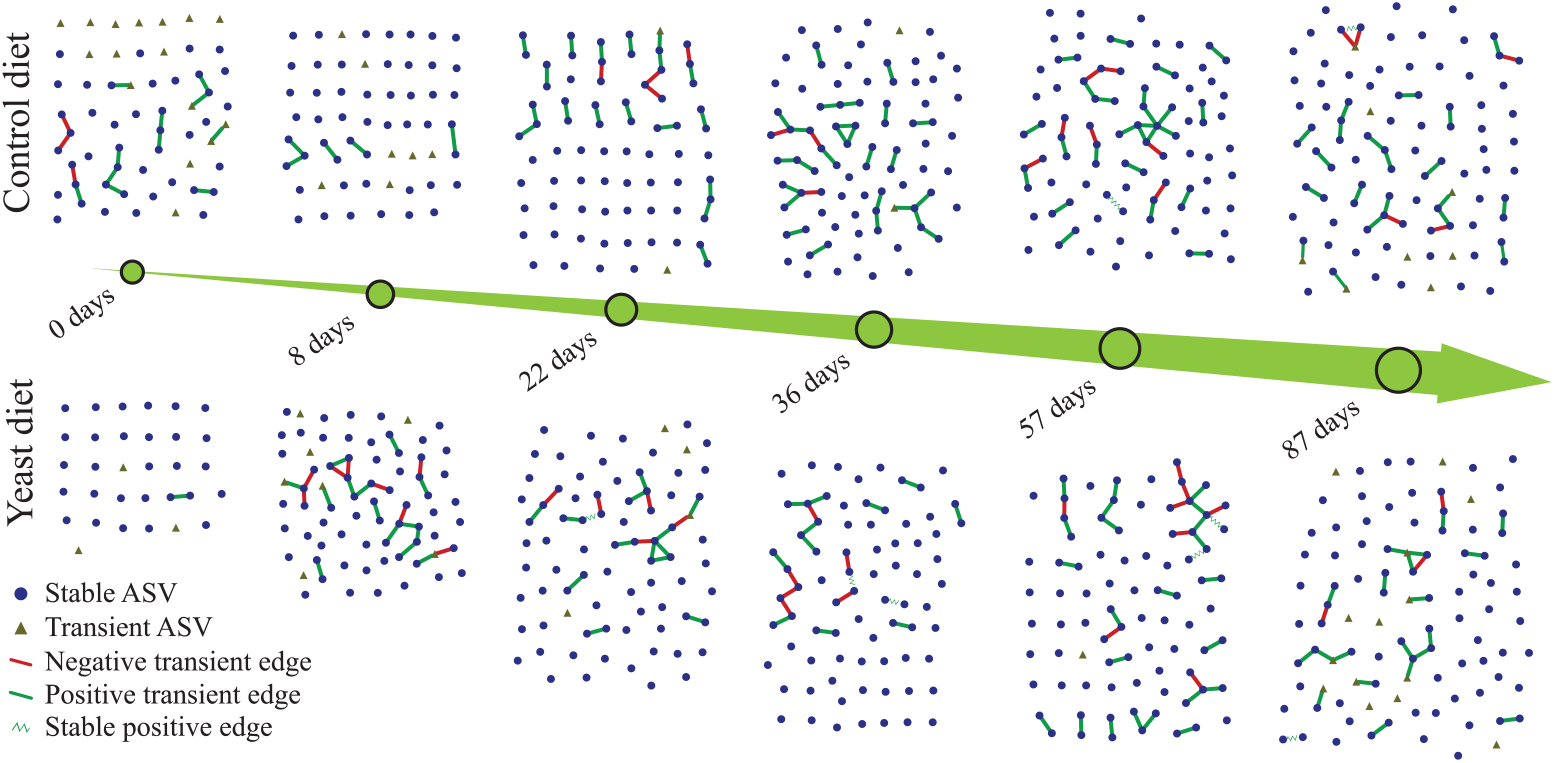
Development of faecal microbial networks across time and feeding groups. **Stable ASVs** are defined as those nodes which were present in at least two consecutive networks. **Transient ASVs** are defined as those nodes which were not present in consecutive networks. **Negative transient edges** are defined as the edges that are present in one network, but do not appear in the following network. The “negative” means the presence of an inverse proportional relationship between two nodes (ASVs). **Positive transient edges** are defined as the edges that are present in one network, but do not appear in the following network. The “positive” means the presence of a proportional relationship between two nodes (ASVs). **Stable positive edges** are defined as the edges that are present in one network and in the following network. The “positive” means the presence of a proportional relationship between two nodes (ASVs)

#### Vertex persistence

There were 11 more ASVs in the YL microbial networks (66 ASVs) that transited throughout the whole experiment compared with those of the CL microbial networks (55 ASVs) (d0 excluded). Those 11 ASVs belonged to 8 different bacterial phyla: *Actinobacteria*, *Bacteroidetes*, *Firmicutes*, *Lentisphaerae*, *Proteobacteria*, *Synergistetes*, *Tenericutes*, and TM7. More specifically, a Coriobacteriaceae family ASV, *Bacteroides* ASV, *Turicibacter* ASV, Peptostreptococcaceae family ASV, *Eubacterium biforme* ASV, and R4-45B, Desulfovibrionaceae, Dethiosulfovibrionaceae, Mycoplasmataceae familiy ASVs transited across the microbial networks starting from d8 PW in the YL faecal microbiomes. In the CL microbiomes, in turn, the transition of a *Blautia* ASV was observed 4 times compared to 3 transitions of that in the YL faecal microbiomes.

#### Edge persistence

When looking at which bacterial genera maintained the same microbe-microbe relationship in more than one consecutive network within diet groups (see methods), there were 3 pairs of ASVs which did so in the YL microbiomes in contrast to one ASV pair in the CL microbiomes. The latter pair of nodes, namely *Asteroleplasma anaerobium* and *Eubacterium biforme* ASVs, were connected at d57 and d87 PW. For the same time points another pair, *Dialister* and *Sutterella*, maintained their connection in the networks recovered from YL microbiomes. The connection between the nodes affiliated to *Prevotella copri* and *Faecalibacterium prausnitzii* ASVs in the YL microbial network was maintained at d22 and d36 PW, the period of transition from weaning to grower-finisher diet. A node representing an unknown ASV of *Clostridiales* order and the node affiliate to *Roseburia faecis* ASV were connected at both d36 and d57 PW.

## Discussion

In this study we attempted to close the knowledge gap on how the gut microbiota develops over time in pigs fed diets in which the SBM/conventional proteins are replaced by yeast-derived proteins during weaning followed by an RSM-based diet during the G/F period. We specifically looked into the possible carry-over effects of the changes in the weaning period faecal microbiomes onto the G/F period microbiomes. As expected, we found differences in alpha and beta microbial diversity between the faecal microbiomes of the yeast-based and SBM-based weaning diets.

We found that the bacterial diversity was higher in the yeast-rapeseed meal (YL) group during the weaning period, which is interesting and contrasts with our previous observations[10] that revealed lower bacterial diversity in caecum and colon microbiomes of the pigs fed with the yeast weaning diet. There are a number of differences between the studies to explain this discrepancy such as: 1) sequencing platform (Illumina Miseq (this study) vs. Illumina Hiseq in [10]; 2) sequencing depth (here we set a threshold of 40000 sequencing reads per sample); 3) 16S *rRNA* gene amplified region (V3-V4 and V1-V3, respectively)[15]; and 4) gastrointestinal (GI) tract originating the samples[15]. In an unpublished study of ours, wherein the caecum and colon microbiomes of pigs challenged with an ETEC *E. coli*, we also observed lower figures of alpha diversity in the microbiomes of pig fed the yeast diet. There too the V3-V4 region of the 16S *rRNA* gene was amplified and sequenced using the Illumina Miseq platform. This suggests that the GI tract region was the factor that contributed the most to the faecal microbiome diversity of the YL piglets. The resident microbial community in caecum or spiral colon can have different structure compared with those of the rectal part because of differences in substrate availability[16]. The way of yeast cell processing used to formulate the yeast-based feed might have had a large impact on the substrate availability in the large intestine hence a distinct microbial community structure. Mannose polymers, the components of the yeast cell wall, cannot be digested by the host[17] and therefore are the substrate for microbial fermentation in the large intestine. *Prevotella* species and *Selenomonas* were shown to be able to degrade mannose among other substrates (summarized in [18]). Our previous results from a study with a nearly identical to this study design showed that the *Prevotella* and *Selenomonas* affiliated taxa were more abundant in the caecum and colon of the weanling pigs fed the yeast-based diet compared with those of the SBM diet[10]. This difference was not replicated in this study in which the faecal microbiomes were analysed. This may suggest that: 1) the microbiome of the proximal part of the large intestine is indeed different than that of the distal part (rectum) and; 2) the bacterial diversity in the yeast related faecal microbiomes is driven by the activity of the bacteria that degrade yeast cell which in turn facilitates the release of nutrients from yeast cells for further bacterial fermentation in the distal part of the colon. To test these hypotheses and to further investigate the microbiome changes due to the yeast-based diets, new research that draws conclusions from metagenomics, transcriptomics, proteomics and other ‘omics’ data at once might be suitable.

Next, we studied the beta-microbial diversity using two methods: unweighted UniFrac[19] which incorporates the phylogenetic information of the microbial communities and weighted UniFrac[20] which incorporates the phylogenetic information of the microbial communities as well as the abundances of the members of the communities. We found that the beta-diversity changes associated with the yeast diet occurred during the weaning period up to d22 PW. It is interesting that the importance of rare microbial species (unweighted UniFrac method) was more pronounced by the end of first week PW than in the end of the weaning period (d22 PW). We hypothesize that the yeast proteins in the feed of weaning piglets is involved in shaping the faecal microbiomes in a two-stage mechanism. First, phylogenetically more distant microbial species establish themselves at low numbers in the yeast related faecal microbiomes by the end of the first week after introduction of the yeast feed. And second, as the yeast feeding lasts for three weeks after the feed introduction, those phylogenetically distinct species increase in numbers, hence making the yeast-influenced microbiomes to cluster apart from the control microbiomes as estimated by the abundance-sensitive weighted UniFrac method.

When we looked into the dynamics of the relative abundance changes of the major bacterial phyla, i.e. *Bacteroidetes* and *Firmicutes*, the results showed that the *Bacteroidetes* fraction decreased on average from 75% to 60% as well as the fraction of *Firmicutes* increased from 20 to 30% in the YL faecal microbiomes during the weaning period. In the CL microbiomes the relative abundance of *Bacteroidetes* and *Firmicutes* seemingly remained invariable at 70% and 20%, respectively, within the same period of the experimentation (shown in Figure 2C). The key finding of this study is that there were less *Bacteroidetes* and more *Firmicutes* by the end of the weaning period (measured at d22 PW) in the yeast group compared with the control group. Although these differences at the phylum level were not retained during the G/F period, when animals were on the RSM-based diet, there was still less *Prevotella* ASVs of *Bacteroides* phylum in the yeast group faecal microbiomes compared with those of the control group. Following up the differentially abundant *Prevotella* in the control group, we discovered a co-occurrence pattern between the *Prevotella* ASV and a *Desulfivibrio* ASV (data not shown). *Desulfivibrio* is a sulphate-reducing hydrogenotrophic species of the pig intestines that participates in hydrogen removing and fermentation[21, 22]. It is intriguing that such a co-occurrence pattern was discovered only in the microbiomes of the control group weanling pigs but not in the yeast piglets. At the later stages of the G/F period (d57, 87 PW) there were differences in low abundance taxa such as *Novispirillum* and *Campylobacter lanienae* between the YL and CL faecal microbiomes. Both bacterial phylotypes represented a small fraction of the faecal microbiomes amounting for less than 1% of all bacterial faecal microbiota. While the presence and function of *Novispirillum* in a pig gut microbiome is less clear, the *C. lanienae* was isolated from faeces of healthy pigs and considered a commensal[23]. Another interesting aspect of finding *C. lanienae* in the faecal microbiomes of pigs fed yeast-based diet during weaning period is that there was a link between *C. lanienae* and the RF3 family of the *Tenericutes* phylum when as per the inference of the respective microbial network.

In order to explore microbe-microbe interactions in the microbiomes of YL and CL, we conducted network analysis by recovering interactions between the ASVs with the SPIEC-EASI algorithm[24]. There were more taxa that were recovered at consecutive time points from the YL microbial networks than that of those of the CL. It means that the members of the YL microbiome, once established during the first week PW, were maintained in the microbial networks until the final phase of G/F period. This also suggests that a combination of yeast diet during weaning and RSM during the G/F period supports the expansion of the core faecal microbiota over those with the control diet during weaning period. Here we apply the term “core microbiome” to designate those bacterial species that are recovered from the faecal samples throughout the whole experiment. Of note, the fraction of the core microbiome to its “non-core” part was 17-20% which suggests that the microbial communities changed dramatically over the period of the experiment. Since our experiment covered nearly the whole life-span of a slaughter pig, it is conceivable that in this study we observed the degree to which the gut microbiome co-evolves together with the host, as seen by the structural changes of the microbiome as a function of the weaning, age, diet, management etc. As stated earlier, the faecal microbial communities may be very different from those that reside in the colon and caecum in terms of their functions and cross-feeding patterns. Our findings, based on the microbial network analysis here, show that only a few taxa were connected at more than one consecutive time points: one pair in the CL microbiomes and three pairs in the YL microbiomes. This suggests that the bacterial interactions were volatile throughout the experiment. Also, the intervals between the sampling events were long enough for the faecal microbiomes to undergo compositional changes hence the possibility of changing the way the microbes interact with each other.

On the other hand, from an ecological perspective, it is within reason for the microbial communities that reside in the terminal part of GI tract where carbohydrate substrate availability is scarce, to switch from an active fermentation to a “hibernation” state on their way out of the habitat. This hypothesis can be tested by analysing microbial networks recovered from samples from both the proximal part of the large intestine (e.g. caecum, colon) and its distal part (rectum) collected post-mortem which was unattainable for this longitudinal study. Previous works investigating pig faecal microbiomes using graph theory methods [25, 26] relied on inferring microbial networks from 16S *rRNA* gene sequencing data using correlation-based approaches[27, 28]. For instance, Kiros and co-workers were able to recover hub bacterial genera having more than 10 connections to other genera of the network when investigating *Saccharomyces cerevisiae* yeast supplementation to weanling piglets, (e.g. *Lactobacillus, Roseburia, Faecalibacterium, Prevotella* etc.) using the CoNet tool for the microbial networks’ recovery[26]. Wang et al., studying pig faecal microbial networks longitudinally by using the SparCC tool, identified more than 10 edges for *Prevotella copri, Blautia, Bacteroides*, and *Faecalibacterium*[29]. In contrast to the mentioned studies, we recovered microbial networks wherein the nodes had 1 connection, or edge, mostly with only few having 3-5 connections. The difference in methodology and possibly a small sample size in this study[24] might have been the non-biological explanation of why the recovered microbial networks were of lower complexity compared to the ones discussed in Kiros et al and Wang et al. Yet, an interesting finding derived from the network analysis was that some pairs of bacterial phylotypes (connected one to another nodes) were observed across several consecutive time points. For instance, in the CL microbiomes, a *Clostridiales/Roseburia faecis*, and *Dialister/Sutturella* bacterial phylotype pairs were recovered in pairs from d36-57 PW and d57-87 PW networks of the G/F period, respectively. Another phylotype pair, *Prevotella copri/Faecalibacterium prausnitzii* was seen connected in both d22 and d36 PW microbial networks. It is intriguing that this finding supports previously discussed carry-over of *Bacteroides* phylum (*Prevotella*) relative abundance from the weaning period onto the beginning of the G/F period. Only one bacterial phylotype pair, *Asteroleplasma anaerobium/Eubacterium biforme*, transited across several time points (d57-d87 PW). This difference in the number of bacterial phylotypes observed across several consecutive time points, can be interpreted as an element of stability that was observed more often in the microbiomes of yeast fed weanling pigs than in that of controls. Also, this type of information may be indicative of the presence of continuous diet-dependent microbe-microbe cross-feeding patterns that is stably expressed during the gut microbiome development.

In conclusion, upon the longitudinal analysis of the pig faecal microbiomes of pigs fed either yeast-based or SBM-based weanling diets, the major differences in the microbiome composition were observed during the second-to-third week post-weaning. Those changes attributed to the differences in the dietary regimes were carried over to the G/F period and primarily represented as a retention of lower relative abundance of *Prevotella* in the yeast microbiomes compared with the control ones; and in the form of the microbe-microbe interactions. To further gain insight into the details of the effect of the animal diets produced in a sustainable way on the gut microbiome of pigs, a study with the exploration of the full genetic context of the entirety of gut microorganisms, that is a collection of all non-host genes, would be of a potential interest. This study seems to support the possible beneficial effect of introducing yeast-based feed ingredients in weanling pigs coupled with the RSM-based feed in the G/F period. The combination of the two sustainably produced feed worked well together rendering a more optimal large intestinal microbiota.

## Materials and Methods

### Ethics statement

The experiment was carried out at the Center for livestock production (SHF) (NMBU, Ås, Norway) approved by the National Animal Research Authority (permit no. 174). All animals were cared for according to laws and regulations controlling experiments with live animals in Norway (the Animal Protection Act of December 20th, 1974, and the Animal Protection Ordinance concerning experiments with animals of January 15th, 1996).

### Animals, allotment, and housing

A total of 48 Norwegian crossbreed pigs (Landrace x Yorkshire x Duroc) from 5 litters were used for the animal performance part of the experiment. Average initial weight and final weight in the piglet period was 10.4 kg and 22.8 kg, and average initial weight and final weight in the growing-finishing period was 22.8 kg and 109.0 kg, respectively. The experiment was conducted as a randomized complete block design. At the start of the piglet period the pigs were blocked by litter and sex and allotted by initial weight to four dietary treatments (below). Piglets were kept in pens with four pigs per pen, giving three replicates per treatment. Each pen had partially slatted floors, and a total area of 2.6 m^2^ (2.6 × 1.0 m). The pens were equipped with heating lamp. A rubber mat of approximately 90 × 100 cm was used as a replacement for other bedding materials, to minimize interference with the measurements of microbiome. The room temperature was kept on average at 19.9°C ± 1.05 SD, with 8 h of light and 16-h darkness cycles. The piglet period lasted 28 days. The piglets were fed *ad libitum* from automatic feeders and had free access to drinking water. After the piglet period, the pigs were moved from the nursery room to a growing-finishing room and re-grouped. The growing-finishing period lasted on average for 89.5 days. At each feeding, pigs were individually restrained in the feeding stall until the feed was consumed in order to obtain individual feed intake. Thus, each pig was one experimental unit. Pigs were housed in an environmentally controlled barn with partially slotted concrete floor. Twelve 8.2 m^2^ pens designed for individual feeding were used. Average ambient daily temperature in the growing-finishing room was 18.5°C ± 1.45 SD.

### Diets and feeding

The dietary treatments in the piglet period were: 1) a control diet based on soybean meal, fish meal, potato protein concentrate and rapeseed meal as protein sources (Control piglet diet), and 2) an experimental diet where 40% of the protein was replaced by *C. jadinii* protein (Yeast piglet diet). After the piglet period, pigs were switched to growing-finishing diets consisting of: 1) a soybean meal based control diet (Control G/F-diet), and 2) a rapeseed meal and field bean based experimental diet (Local G/F diet). The diets were designed to be isonitrogenous and isoenergetic and to contain equal levels of methionine + cysteine, and threonine. The diets were produced and pelleted to 3 mm diameter at a commercial feed factory (Felleskjøpet Kambo, Moss, Norway). The content of digestible lysine, threonine, methionine and cysteine of the ingredients was estimated using analyzed values, multiplied by the standardized ileal digestibility coefficients (SID) for nitrogen and amino acids[30]. All diets were formulated to meet or exceed the requirements for indispensable amino acids and all other nutrients[31]. A cumulative feed sample from each dietary treatment was taken for chemical analysis. Composition and nutrient contents of diets are shown in Table S3 (piglet diets) and Table S4 (G/F diets)[32]. When combining the piglet period and the G/F period, the following four treatments were obtained: 1) Piglet control diet + G/F control diet. (Control/Control, or CC) 2) Piglet control diet + G/F local diet. (Control/Local, or CL) 3) Piglet yeast diet + G/F control diet. (Yeast/Control, or YC) 4) Piglet yeast diet + G/F local diet (Yeast/Local, or YL)

In the piglet period the pigs were fed pen-wise according to appetite. All four pigs in each pen were given the same feed. The average weight gain and feed intake for each pen was measured weekly, and average daily gain (ADG), average daily feed intake (ADFI) and feed conversion rate (FCR) as kg feed divided on kg gain were calculated for each pen.

In the growing-finishing period (G/F period), all pigs were individually fed twice per day according to a semi-ad libitum feeding scale[33]. Feed refusals for each pig were recorded and subtracted from the total feed intake. All pigs were given free access to water from nipple drinkers. Feed consumption and individual pigs’ weight were recorded weekly to determine average daily gain (ADG), average daily feed intake (ADFI) and FCR.

### Chemical analyses

Samples of the diets were analysed for crude protein (CP) by Kjeldahl-N x 6.25 (EC No 152/2009), crude fat using ASE^®^ 350 Accelerated Solvent Extractor, dry matter (DM) by drying to constant weight at 104°C (EC No 152/2009), ash by incineration at 550°C (EC No 152/2009), acid detergent fiber (ADF) and neutral detergent fibre (NDF) using a fibre analyser system (Ankom200; ANKOM Technologies, Fairport, NY, USA) with filter bags (Ankom F58; ANKOM Technologies). Gross energy (GE) content was determined by a Parr 1281 Adiabatic Bomb Calorimeter (Parr Instruments, Moline, IL, USA) according to ISO (1998). Analysis of amino acids in the diets were carried out according to EC (2009) using Biochrom 30 Amino Acid Analyzer. Tryptophan in the diets was determined according to EC (2009) using high-performance liquid chromatography system (Dionex UltiMate 3000, Dionex Softron GmbH, Germering, Germany) and the fluorescent detector (Shimadzu RF-535; Shimadzu Corp., Kyoto, Japan).

### Faecal sample handling

Faecal samples were collected from 8 pigs of CL and 8 pigs of YL group chosen randomly from the animals involved in the animal performance part of the experiment (see Animals, allotment, and housing). The collection of faecal samples was carried out at d0, d8, d22, d36, d57, and d87 post-weaning (PW). On d87 PW there were 7 samples from each group. The samples were liquid nitrogen snap frozen and kept at −80°C until the DNA isolation. The DNA extraction was according to a previously described protocol[34] with minor modifications. Briefly, 200 mg of thawed and mixed faecal samples were added to 1 ml of InhibitEX Buffer (QIAGEN, GmbH, Hilden, Germany) followed by the beat-beating step in TissueLyser II (Qiagen, Retsch GmbH, Hannover, Germany) with 500 mg of zirconia/silica beads (Ø = 0.1 mm, Carl Roth, Karlsruhe, Germany) (1.5 min at 30 Hz). Proteins were digested with 30 *μ*L of Proteinase K II (QIAGEN, GmbH, Hilden, Germany). DNA was washed with AW1 and AW2 buffers (QIAGEN, GmbH, Hilden, Germany) and eluted with ATE buffer (QIAGEN, GmbH, Hilden, Germany). The yielded DNA purity was assessed by NanoDrop (Thermo Fisher Scientific, Waltham, MA) and quantified with the Qubit fluorometric broad range assay (Invitrogen, Eugene, OR, USA). Library preparation was performed at the Norwegian Sequencing Centre (https://www.sequencing.uio.no/, Oslo, Norway) using universal prokaryotic primers 319F(5’-ACTCCTACGGGAGGCAGCAG-3’) and 806R(5’-GGACTACNVGGGTWTCTAAT-3’) that target the V3-V4 regions of the 16S *rRNA* gene. Sequencing was performed on a MiSeq sequencer following the manufacturer’s guidelines. The resulting sequences were deposited in the ENA (PRJEB41040). Metadata can be accessed through https://github.com/stan-iakhno/bioRxiv_02.

### Bioinformatics analysis and statistics

Demultiplexed paired-end Illumina reads were pre-filtered with bbduk version 37.48 (BBMap – Bushnell B., https://sourceforge.net/projects/bbmap/) by trimming right-end bases less than 15 Phred quality score, removing trimmed reads shorter than 250 bp or/and average Phred quality score less than 20. The resulting reads were further quality filtered by trimming left-end 20 bp and removing reads with maxEE more than 1 for forward and 2 for reverse reads, denoised, merged, and chimera removed with DADA2 R package ver 1.12.1[35] (Figure S1). The resulting ASV tables that derived from two separate Illumina sequencing runs were merged followed by taxonomy assignment using RDP Naive Bayesian Classifier implementation in DADA2 R package (default settings) with GreenGenes database version 13.8 [36] as the reference database. The phylogenetic tree was reconstructed under the Jukes-Cantor (JC) nucleotide model with gamma distribution (number of intervals of the discrete gamma distribution (k)=4, shape=1 with invariant sites (inv=0.2)) in R. The pipeline code is available through https://github.com/stan-iakhno/bioRxiv_02.

DivNet statistical procedure[37] was used to estimate the Shannon diversity index and to test for differences in Shannon diversity estimates in networked gut microbial communities stratified by the day of sampling with the diet as a covariate. The beta diversity analysis was performed via the analysis of multivariate homogeneity of group dispersions[38] followed by the permutation test[39], 9999 permutations and principle coordinate analysis (PCoA) on unweighted[19] and weighted[20] Unifrac distances, and permutational multivariate analysis of variance (PERMANOVA) test for covariate significance in R, 9999 permutations. The samples with read count less than 40000 were discarded from the alpha and beta diversity analyses. To calculate the relative abundance of bacterial phylotypes per feeding group and per sampling time point, the group means were taken from the respective groups. To detect differentially abundant bacterial phylotypes, ‘corncob’ algorithm[40] was run on the microbial feature tables (ASV counts per each sample) by fitting a beta-binomial regression model to microbial data stratified by the day of sampling with the diet and litter as covariates. The false discovery rate due to multiple testing was addressed by the Benjamini-Hochberg correction with the cut-off of 0.05. The test was run at each taxonomic level (phylum, class, order, family, species, and ASVs) discarding the samples with the read count less than 10000. Those ASVs that lacked genus/species taxonomic classification, were classified manually by using web-based nucleotide BLAST on the non-redundant nucleotide database where possible. Ambiguous hits were ignored.

### Microbial network analysis

The ASV counts were collapsed at the genus level and filtered for at least 3 counts per ASV in at least 20% of the samples and at least 50% of the sample per time point (0, 8, 22, 36, 57 and 87 days) and condition (yeast diet and control diet) using the R package phyloseq[41] version 1.26.1. For each time point and condition a network was computed with the package SpiecEasi[24] version 1.0.7. For each condition the permanence of nodes (ASVs) and edges (their relationships) was checked at two consecutive time points.

## Acknowledgments

We thank the staff at the Centre for livestock production (SHF), NMBU, Ås, for their excellence in managing the animals during this experiment. SI was funded by a PhD fellowship from the Department of Food Safety and Infection Biology at Norwegian University of Life Sciences. The authors thank Foods of Norway Centre for Research-based Innovation (The Research Council of Norway, Lysaker, Norway, grant number 237841/030) and the Centre’s industrial partners for financial support.

## Supporting Information

**Table S1.** Performance results from weaning until slaughter. For the overall experimental period (from average live weight of 10.4 kg until slaughter) no significant differences among four treatments, control-control(CC), control-local(CL), yeast-control(YC),and yeast-local(YL), were found for average daily gain (ADG) (P=0.671) and average daily feed intake (ADFI) (P=0.924). Feed conversion rate (FCR) was influenced by treatment (P=0.032), and pigs given the control diet (CC and YC) in the growing-finishing (G/F) period in general had better FCR than the pigs fed the Local diet (CL and YL). In the piglet period (live weight 10.4 kg until 22.8 kg), FCR did not differ among treatments (P=0.994). Different letters indicate significant difference among treatments (P < 0.05). Average daily gain - ADG. Average daily feed intake – ADFI, and Feed conversion ratio – FCR

|  | CC | CL | YC | YL | Std.err | P-value |
| --- | --- | --- | --- | --- | --- | --- |
| No pens | 3 | 3 | 3 | 3 | - | - |
| Pig per treatment | 12 | 12 | 12 | 12 | - | - |
| Initial weight, kg | 10.37 | 10.32 | 10.44 | 10.37 | 0.731 | 0.9995 |
| Weight after 4 week | 22.84 | 22.80 | 22.54 | 23.11 | 1.092 | 0.986 |
| Final weight, kg | 108.87 | 109.78 | 109.97 | 107.43 | 1.597 | 0.676 |
| Overall ADG, g | 841.4 | 837.5 | 853.7 | 829.8 | 13.62 | 0.671 |
| Overall ADFI, g | 1638.8 | 1653.8 | 1629.8 | 1652.5 | 29.23 | 0.924 |
| Overall FCR | 1.948ab | 1.974b | 1.909a | 1.991b | 0.016 | 0.032 |

**S1 Figure.**
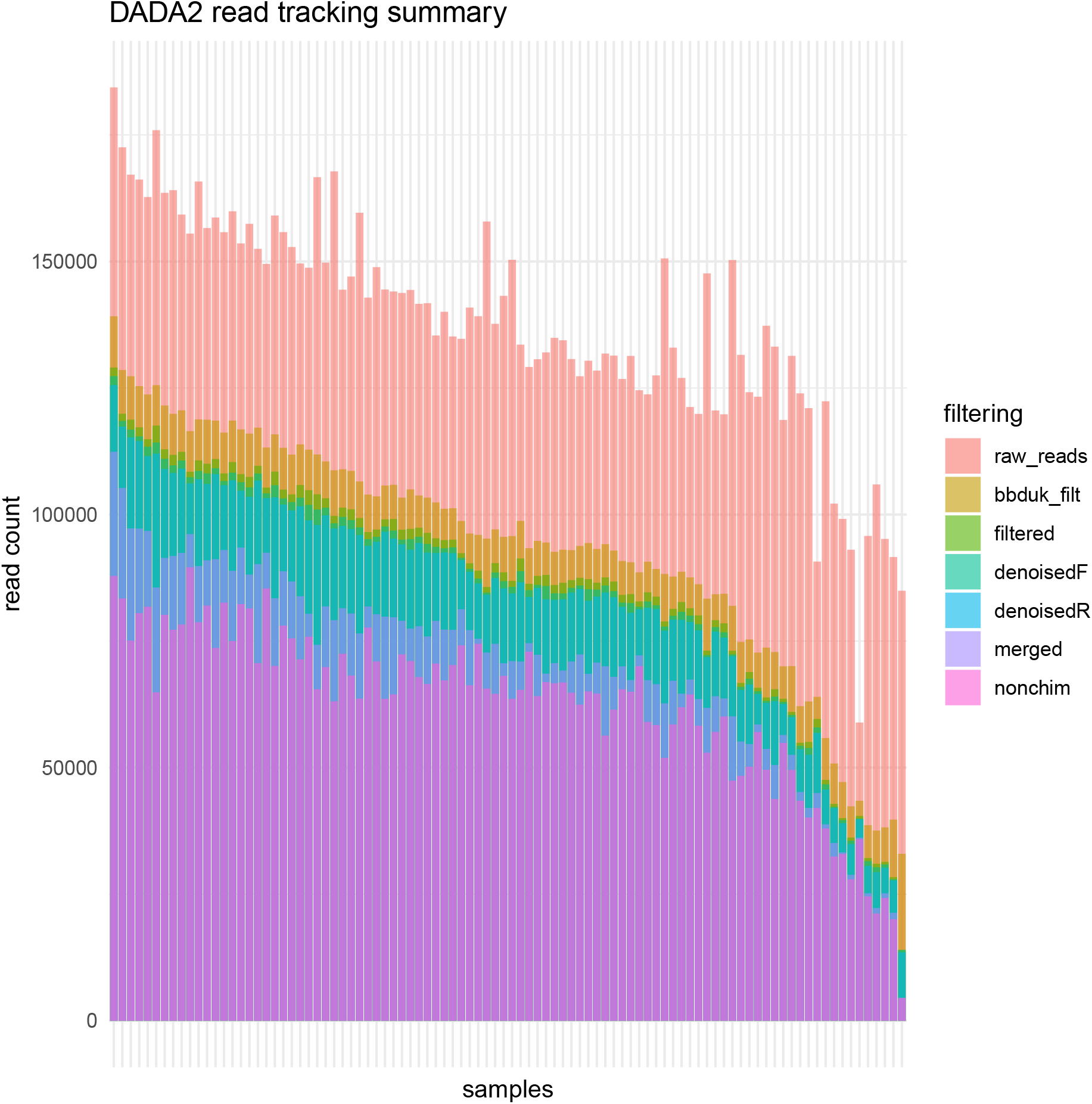
Read tracking summary. The bottom-most bars in the stack (**nonchim**) show the number of read that were the basis for making the feature count table (OTU/ASV-table). The bars above **nonchim** summarise the number of sequencing reads removed due to each procedure of the bioinformatics pipeline: a) filtered with the bbduk filtering algorithm (**bbduk filt**), b) filtered with the DADA2 algorithm (**filtered**), c) removed due to DADA2 denoising procedure (**denoisedR/F**), d) removed due to pair merging failures (**merged**). **raw reads** are raw demultiplexed reads derived from Illumina sequencer.

**S2 Table.** Beta diversity PERMANOVA and permdisp test. The tests were performed on the “Unifrac” and “weighed Unifrac” distances. The test statistics of the permutational multivariate analysis of variance (PERMANOVA) test are given in normal font, multivariate homogeneity of groups dispersions (permdisp) test are given in **bold**.

| *Unifrac* |  |  |  |  |  |  |
| --- | --- | --- | --- | --- | --- | --- |
| *day PW->* | *0* | *8* | *22* | *36* | *57* | *87* |
| *F* | *1.508* | *1.704* | *1.165* | *0.844* | *1.033* | *1.26* |
| *R<sup>2</sup>* | *0.143* | *0.146* | *0.077* | *0.057* | *0.074* | *0.095* |
| *p-value* | *0.125* | *0.024* | *0.212* | *0.704* | *0.359* | *0.122* |
| ***F*** | ***0.002*** | ***0.444*** | ***0.113*** | ***2.57*** | ***2.133*** | ***0.049*** |
| ***p-value*** | ***0.976*** | ***0.527*** | ***0.748*** | ***0.122*** | ***0.168*** | ***0.83*** |
| *weighted Unifrac* |  |  |  |  |  |  |
| *day PW->* | *0* | *8* | *22* | *36* | *57* | *87* |
| *F* | *2.031* | *0.916* | *4.893* | *1.583* | *0.492* | *1.927* |
| *R<sup>2</sup>* | *0.184* | *0.084* | *0.259* | *0.102* | *0.036* | *0.138* |
| *p-value* | *0.104* | *0.491* | *0.003* | *0.156* | *0.821* | *0.115* |
| ***F*** | ***1.013*** | ***0.41*** | ***1.59*** | ***3.302*** | ***0.944*** | ***0.969*** |
| ***p-value*** | ***0.364*** | ***0.543*** | ***0.258*** | ***0.092*** | ***0.354*** | ***0.328*** |

**S2 Figure.**
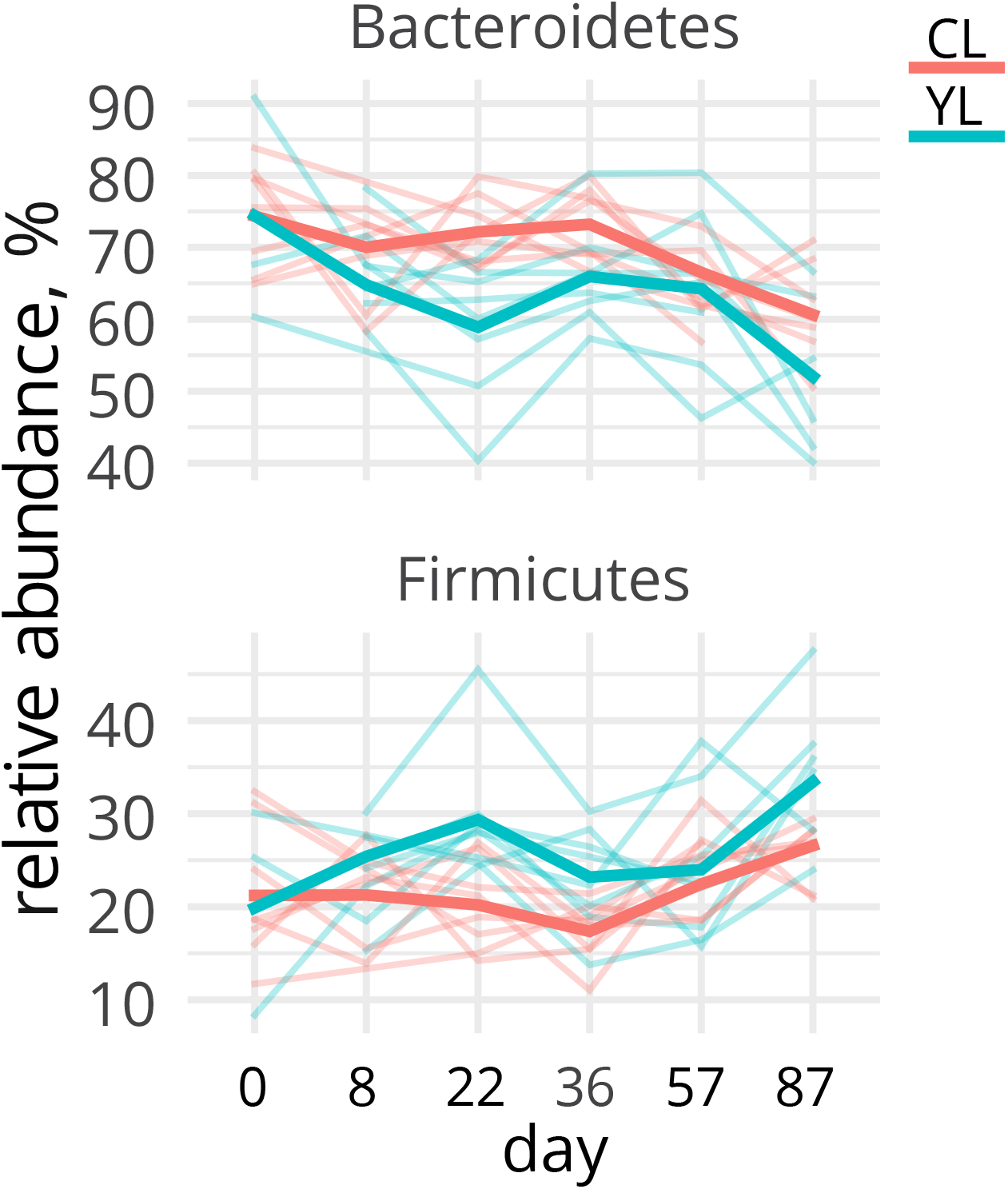
Relative abundance of *Bacteroidetes* and *Firmicutes* phyla across d0-88 PW. Individual observations are shown by the thin spaghetti lines, the average group values are shown by the thick spaghetti lines

**Table S3:**
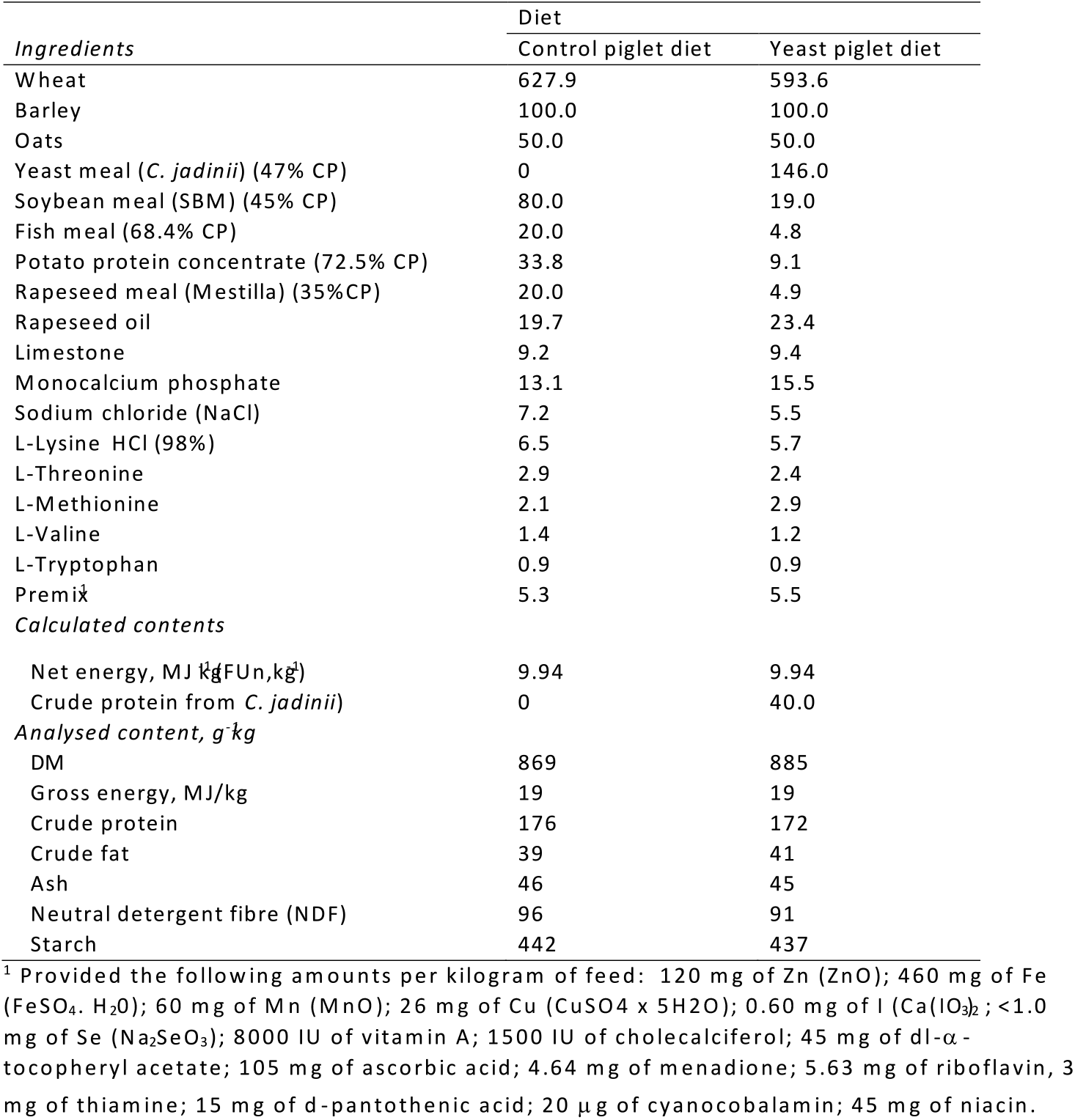
Piglet period. Ingredient and chemical composition (g kg^−^1) of diets based on soybean meal (Control) and *C. jadinii* (Yeast). In the yeast diet, 40% of the crude protein was replaced by that from *C. jadinii* (LYCC-7549; Lallemand Yeast Culture Collection).

**Table S4:** Growing-finishing period. Ingredient and chemical composition (g kg^−^1) of diets based on soybean meal (Control) and locally produced protein ingredients (Local).

| *Ingredients* | Diet |  |
| --- | --- | --- |
|  | Control G/F diet | Local G/F diet |
| Barley | 585.5 | 456.2 |
| Oats | 150.0 | 150.0 |
| Soybean meal (SBM) (45% CP) | 142.6 | 0.0 |
| Rapeseed meal (RSM) (Mestilla) | 60.0 | 180.0 |
| Faba beans (Columbo) | 0.0 | 161.12 |
| Rendered fat | 23.0 | 16.5 |
| Molasses | 10.0 | 10.0 |
| Limestone | 9.7 | 8.7 |
| Monocalcium phosphate | 3.2 | 1.4 |
| Sodium chloride (NaCl) | 6.3 | 6.4 |
| L-LysineHCl (98%) | 3.3 | 3.0 |
| L-Threonine | 1.3 | 1.4 |
| L-Methionine | 1.0 | 1.0 |
| L-Tryptophan | 0.1 | 0.3 |
| Premix | 4.0 | 4.0 |
| *Calculated contents* |  |  |
| Net energy, MJ/kg FUn, kg<sup>1</sup> | 9.42 (1.07) | 9.42 (1.07) |
| *Analyzed content, g/kg* |  |  |
| DM | 868 | 870 |
| Gross energy, MJ/kg | 16.4 | 16.6 |
| Crude protein | 153 | 158 |
| Crude fat | 40 | 51 |
| Ash | 45 | 44 |
| Neutral detergent fiber (NDF) | 165 | 173 |
| Starch | 424 | 422 |

<sup>1</sup> Provided the following amounts per kilogram of feed: 20 mg of Aextra phytase, 72 mg of Zn (ZnO); 96 mg of Fe (FeSO<sub>4</sub> · H<sub>2</sub>O); 48 mg of Mn (MnO); 17 mg of Cu (CuSO<sub>4</sub> x 5H<sub>2</sub>O); 0.48 mg of I (Ca(IO<sub>3</sub>)<sub>2</sub>); 0.27 mg of Se (Na<sub>2</sub>SeO<sub>3</sub>); 6500 IU of vitamin A; 1500 IU of cholecalciferol; 75 mg of dl- $\alpha$ -tocopheryl acetate, 150 mg of Vitamin E (50%); 4.63 mg of menadione; 5.625 mg of riboflavin, 15 mg of d-pantothenic acid; 15 $\mu$ g of cyanocobalamine; 45 mg of niacin; 0.30 mg of biotin; 1.69 mg of folic acid; Choline: 2300 mg (Control) and 1605 mg (Local).

